# A Nested 2-Level Cross-validation Ensemble Learning Pipeline Suggests a Negative Pressure Against Crosstalk snoRNA-mRNA Interactions in *Saccharomyces Cerevisae*

**DOI:** 10.1101/293555

**Authors:** Antoine Soulé, Jean-Marc Steyaert, Jéerôme Waldispuühl

## Abstract

The growing number of RNA-mediated regulation mechanisms identified in the last decades suggests a widespread impact of RNA-RNA interactions. The efficiency of the regulation relies on highly specific and coordinated interactions, while simultaneously repressing the formation of opportunistic complexes. However, the analysis of RNA interactomes is highly challenging due to the large number of potential partners, discrepancy of the size of RNA families, and the inherent noise in interaction predictions.

We designed a recursive 2-step cross-validation pipeline to capture the specificity of ncRNA-mRNA interactomes. Our method has been designed to detect significant loss or gain of specificity between ncRNA-mRNA interaction profiles. Applied to snoRNA-mRNA in Saccharomyces Cerevisae, our results suggest the existence of a repression of ncRNA affinities with mRNAs, and thus the existence of an evolutionary pressure inhibiting such interactions.

## 1 Introduction

Evidence of the breadth of the role of ribonucleic acids in gene regulation are now multiplying. For instance, in eukaryotes microRNAs bind mRNAs to control gene expression [1], and in prokaryotes the OxyS RNA interacts with the fhlA mRNA to prevent ribosome binding and thus inhibit translation [2].

Among all non-coding RNAs (ncRNAs) already identified, the category of small nucleolar RNAs (snoRNAs) is of particular interest. snoRNAs form a large class of well-conserved small ncRNAs that are primarily associated with chemical modifications in ribosomal RNAs (rRNAs) [3]. Recent studies revealed that orphan snoRNAs can also target messenger RNAs (mRNAs) in humans [4] and mice [5], and probably contribute to regulate expression levels. However, despite recent investigations there are to date no evidence that similar snoRNA-mRNA interactions occur in simpler unicellular microorganisms [6, 7].

Interestingly, it turns out that RNA-based gene regulation mechanisms have been primarily linked to higher eukaryotes [8], although it is still not clear if this observation results from an incomplete view of RNA functional landscape or the existence of a negative pressure preventing RNA to interfere with other transcripts.

Our understanding of RNA-mediated regulation mechanisms significantly improved in recent years. In addition to well-documented molecular pathways (e.g. [2]), regulation can also occur at a higher level through global affinities between ncRNAs and mRNAs populations [9]. Furthermore, Umu *et al*. [10] showed another intriguing, yet complementary, level of control of gene expression that could explain discrepancies previously observed between expressions of mRNAs and the corresponding protein expressions in bacteria [11, 12]. In their study, the researchers extracted a signal suggesting a negative evolutionary pressure against random interactions between ncRNAs and mRNAs that could reduce translation efficiency. However, these results cannot be trivially extended to eukaryotes where the role of the nucleus has to be considered.

In this study, we investigate this phenomenon of avoidance of random interactions between ncRNA and mRNA in *Saccharomyces Cerevisae*. In particular, we focus our analysis on the bipartite interactome between snoRNAs and mRNAs. Indeed, the snoRNA family is an ancient and large class of ncRNAs for which the mechanism of mRNA avoidance could explain the absence of known interactions between snoRNAs and mRNAs in unicellular eukaryotes.

A major challenge of this analysis stems from severely unbalanced datasets. While we retrieve more than 6000 annotated mRNAs, we could only recover less than one hundred snoRNAs [13]. Such disparity is a serious source of bias that should be carefully addressed. Therefore, we developed a customized ensemble learning pipeline to quantify the specificity of RNA binding profile between unbalanced RNA families.

First, we use state-of-the-art prediction tools to compute the snoRNA-mRNA interactome as the set of all interactions between snoRNAs and mRNAs. Then, we design an ensemble learning pipeline to identify statistically significant biases in the distribution of binding affinities between classes of RNAs. Importantly, in order to remove any possible source of bias during the parametrization of classifiers, we introduce a second level of Leave-One-Out Cross-Validation (LOOCV) to avoid overfitting. Our results reveal that although classes of snoRNAs exhibit preferential interaction patterns with mRNAs, this selective pressure is not as strong as initially anticipated. It corroborates previous hypothesis on prokaryotes, and suggests the presence of a phenomenon of avoidance of random interactions between ncRNAs and mRNAs in single-celled eukaryotes.

## 2 Approach

We aim to characterize the strength and specificity of random ncRNA-mRNA interactions in *Saccharomyces Cerevisae*, although our work primarily focuses on snoRNA-mRNA interactions. Our data set includes smaller categories of ncRNAs (e.g. spliceosomal RNAs) used for an additional control of our results.

We computed ncRNA-mRNA interactomes from ncRNAs and mRNAs sequences using two different state-of-the-art computational prediction tools (RNAup [14] and intaRNA [15]). Those predictions are to serve as an approximation for the propensity of those ncRNAs to form crosstalk interactions with mRNAs. By using an ensemble learning pipeline, we approximated the specificities of interaction profiles in those interactomes. We also approximated the specificities of ncRNAs sequences with machine learning upon the Kmer compositions of the said sequences. The comparison of the approximated specificities highlights a global pressure inhibiting the affinity ncRNA-mRNA interactions in *Saccharomyces Cerevisae*.

We finally completed this work by a collection of complementary control tests providing a better understanding of the limitations of this work. Data, code, raw results and supplementary displays are available at http://jwgitlab.cs.mcgill.ca/Antoine/nested_loocv_pipeline/tree/master.

## 3 Methods

### 3.1 Dataset

#### Saccharomyces Cerevisae

We focus our study on a single organism: Saccharomyces Cerevisae. Working on a single organism ensures that all the molecules co-evolved and that their interactions were under the same evolutionary pressure. We also focus our study on a eukaryote to investigate the influence of the nucleus. Indeed, the nuclear membrane creates a confined environment that segregates molecules. Moreover, eukaryotes usually display more complex mechanisms and have more coding sequences than prokaryotes and archaea. Extending the study to a family instead or even further, like Umu *et al*. [10] did for instance, has been considered. However, less data are available for other related yeasts and including more species increases the number of parameters to consider. We came to the conclusion that a multi-species study, while being interesting, was unrealistic yet. Finally, we excluded multicellular organisms to avoid problematic phenomenons like specialized tissues.

For all those reasons, this study required a unicellular eukaryote offering a satisfying number of identified RNA sequences and *Saccharomyces Cerevisae* appeared to be the most suited model by being a model eukaryote organism with the greatest number of annotated sequences amongst unicellular eukaryotes. All sequences have been obtained from the manualy curated Genolevure [13] database.

#### ncRNA-mRNA Interactome

Our main source of features is the ncRNA-mRNA interactome i.e. all the ncRNA-mRNA interactions. Noticeably, we are referring to ncRNA-mRNA interactions as computational predictions instead of experimentally observed interactions. The probability of such event is conventionally approximated by the energy barrier and difference of entropy (*∆g*) between the structures of the two molecules and the structure of a potential complex [14]. We work under the usual and reasonable assumption that, for two complexes *i* and *j*, if 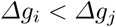 then the complex *i* is more stable than the complex *j* and thus is more likely to form and be observed. In order to study the set of all potential ncRNA-mRNA complexes, we computed for all {ncRNA,mRNA} pairs the corresponding *∆g* using prediction tools (cf. section 3.2), thus resulting in two predicted ncRNA-mRNA interactomes: one for each prediction tool we used (See section 3.2).

We focused our study on ncRNA-mRNA interactome for two reasons. First, the role of mRNA as temporary medium of genetic material makes it a central element in most cellular pathways. mRNAs are centrepieces of several mechanisms such as regulation [10–12] and splicing that might be impacted by crosstalk interactions. Second, ncRNAs (i.e. non-coding RNA, which refers here to RNA which are neither messenger, transfer or ribosomal RNA and also excludes miRNA and siRNA cf. section 3.1) offer properties of interest for this study.

Indeed, the selected ncRNAs can be clustered into categories sharing similar properties, such as structure and length, which makes any comparison more meaningful. Those ncRNAs are also free from cellular mechanisms such as maturation or directed export that might generate noise. Finally, there is no observed interaction between those ncRNAs and mRNAs. As a consequence we can assume than the interaction we predict are opportunistic and not part of a defined biological pathway. A detailed description of ncRNAs labels is provided in section 3.1 and in the supplementary material.

We also considered two other practical aspects in this decision: maximizing the number of available annotated sequences and maximizing the number of crosstalk interactions (i.e. minimizing the number of known interactions). The first aspect directly impacts the statistical validity of any potential results and the second is justified by the goal of this study. The ncRNA-mRNA interactome also satisfies those two aspects.

#### ncRNA Labels

In order to conduct this study, we had to choose which of the mRNAs or ncRNA to label. The absence of structural properties in mRNA naturally inclined us to label ncRNA instead. We produced a 5-label classification (cf. Table 1) according to the gene ontologies based on both functional and structural properties. Out of those five labels, two labels happen to be much more similar in terms of lengths and numbers. As a consequence, we performed all our tests with both the five labels dataset and a dataset limited to those two similar labels and are providing displays for both.

**Table 1.** Numbers for both 5-label and 2-label datasets, means and standard deviations of distributions of sequence lengths and the colours associated in our displays for each ncRNA label

| ncRNA<br>Label | Dataset |  | Length |  | colour |
| --- | --- | --- | --- | --- | --- |
| | 5 labels | 2 labels | $\mu$ | $\sigma$ | |
| miscellaneous | 11 | 0 | 900.27 | 828.04 | black |
| C/D box | 45 | 45 | 106.87 | 34.41 | red |
| H/ACA box | 29 | 29 | 270.83 | 176.75 | blue |
| spliceosomal | 5 | 0 | 245.60 | 183.40 | yellow |
| unknown | 7 | 0 | 500.14 | 225.31 | green |
| total | 97 | 74 | 281.39 | 383.93 |  |

A complete description of all those ncRNA is available in the supplementary material. For the sake of clarity, we will only provide a shorter description of each label in this paper.

C/D box and H/ACA box ncRNAs are snoRNAs (small nucleolar RNA) involved in pre-rRNA maturation by performing two different modifications of specific bases. C/D box snRNAs are performing pseudouridylation, an isomerization of uridines into pseudouridines. Pseudouridines have an extra NH group able to form supplementary hydrogen bonds. Those bonds stabilize rRNA structure [16, p.200]. H/ACA box snRNAs are performing 2-O methylation, a methylation of the ribose. RNA has a short lifespan compared to DNA. By methylating the ribose, the rRNA is less vulnerable to degradation by bases or RNAses. In addition to this increased lifespan, this modification also impacts the rRNA structure by changing spatial constraints and decreasing the number of hydrogen bonds the modified base can form [16, p.200].

Those two labels are the most consistent, both in numbers and internal similarities with both sequential constraints (boxes) and similar structures common to all the ncRNAs of a given label. Moreover, the lengths of ncRNAs are consistent inside each label and shorter than the ncRNA average (cf. Fig 4).

Importantly, we will use these two groups (i.e. C/D box and H/ACA box ncRNAs) to study the existence of an evolutionary pressure on snoRNAs. The other groups described below will be used as control and/or to suggest the generalization of the pressure to other classes of ncRNAs.

Spliceosomal ncRNAs share the common trait of being involved in the splicing process. However all other properties vary.

Miscellaneous ncRNAs have been identified and their functions are known. However those functions are too specific and diverse to be gathered in any label but Miscellaneous. Moreover all other properties vary.

Unknown ncRNAs have been identified but, unlike miscellaneous ncRNAs, their functions remain unknown. Moreover all other properties vary in an even wider range than the two previous labels.

### 3.2 Features Description

This section describes the three metrics used in this study to produce the main sets of features: RNAup, IntaRNA and Kmer composition similarity. Other basic features used as control, such as sequence length, are not described as they are straightforward.

#### RNA-RNA Interactions Prediction Tools

In order to produce a satisfying interactome we use two different RNA-RNA interaction prediction tools: RNAup[14] and IntaRNA[15, 17]. We selected nonspecialized prediction tools over specialized ones such as RNAsnoop [18] as we are interested in non-specific interaction.

Both RNAup and IntaRNA implement the same core strategy. They compute the hybridization energies between the two RNAs as well as the accessibility (i.e. probability of being unpaired) for each interaction site. Those values are then combined to score potential interaction sites. The highest scoring sites are returned together with the free energy of binding. We can then retrieve the secondary structures of each individual RNA using constraint folding algorithms.

RNAup strictly implements this strategy thus predicting the optimal minimum free energy (MFE) compatible with the axioms. IntaRNA differs by two aspects. The first one is that the version of IntaRNA used in this work uses a slightly less recent version of Turner energies model. However the differences between those versions are minor and are very unlikely to produce the observed dissimilarities. The second one is that IntaRNA adds a seeding step to reject interaction sites deemed unlikely. This extra step reduces the search space by focusing on the most promising ones and significantly reduces the runtimes compared to RNAup. An extensive description of the seeding procedure is presented by *Bush et al.* [17]. Comparative benchmarks place IntaRNA in the top of prediction tools with better scores than RNAup [19, 20]. Indeed, IntaRNA appears to predict interactions closer to the observed ones compared to predictions from others prediction tools, including RNAup. As a consequence this heuristic seems well founded and efficient.

In this study we used this difference between RNAup and IntaRNA to predict two slightly different interactions modes. For each {ncRNA,mRNA} pair, we are assuming that RNAup outputs the optimal MFE regardless of its likelihood while IntaRNA outputs a probably weaker but more realistic interaction. Since realistic interactions are more likely to be observed in the cell than the theoretical optimums, any pressure should impact the first before the second. As a consequence, we aimed at highlighting such pressure by studying those two sets of interactions in parallel.

#### Kmer Composition Similarity

In addition to the two prediction tools mentioned in the previous subsection, we use a third metric: the similarity of the Kmer composition of ncRNA sequences. The term *Kmer* refers here to every possible sequence of nucleobasis of length *K*. This metric associates to each ncRNA the distribution of each Kmer in its sequence, including repetitions. The set of all those distributions is gathered as a vector space suitable for machine learning (cf. 3.3). We produced this third set of features in order to assess the specificity of the sequence and to provide a reference point to the two other sets of features.

All experiments involving Kmers have been made with K = 5 for two reasons. The first one is that five is the length of the average interacting zone in RNA-RNA interactions and so is a suitable length to capture any key subsequences impacting those interactions. The second one is that the number of Kmers to consider grows with the value of K. K = 5 offers the advantage of being both manageable in term of cost and also results in a number of dimensions comparable to the two other methods (i.e. RNAup and IntaRNA). We performed preliminary tests with others values, especially K = 6. Those tests showed little to no differences.

### 3.3 Ensemble Learning Pipeline

The overall goal of our machine learning approach is to investigate a possible bias affecting ncRNA-mRNA crosstalk interactions. In order to do so, we compare the specificity of ncRNA sequences with the specificity of ncRNA-mRNA interaction profiles. The specificity of ncRNA sequences is approximated by the ability of classifiers to predict the labels of ncRNAs from their Kmer composition. ncRNA-mRNA interaction profiles are predicted using prediction tools and their specificity is approximated by the ability of classifiers to predict the labels of ncRNAs from those profiles.

Our utilization of machine learning in this project is challenging for two different reasons that justify all the following methodology choices:

1. The ratios *|vectors|/| features|* of our datasets are problematic: 97 vectors for 6663 dimensions for the vector spaces built from the interactomes. Those ratios are due to both cellular biology, since the number of mRNAs in a genome is always greater by several folds to the number of ncRNAs, and the limited availability of annotated ncRNAs sequences thus limiting as well the number of vectors. Those two issues are beyond our control and, to our knowledge, there is no way for us to significantly improve those ratios for Saccharomyces Cerevisae without considerable drawbacks. Moreover Saccharomyces Cerevisae already has the greater number of annotated ncRNAs amongst similar organisms.

2. Our goal is neither to train a good classifier nor to classify unlabelled RNAs but to estimate how well the labels can be predicted from the different features. We are working under the reasonable assumption that a loss in performance between two sets of features implies that the lesser performing set is less specific. If the two sets of features are related, like ours are, it would imply a levelling mechanism.

#### Leave-one-out Cross-validation (LOOCV)

Cross-validation refers in machine learning to partitioning the data set into different sets to separate the data used to train the classifier and the ones used to test it. The goal of cross-validation is to ensure the credibility of the results produced.

We use a leave-one-out cross-validation technique (LOOCV) for validation. For every vector *v_i_* in our set *V* of vectors we train a classifier on the set (*V − v_i_*) and test the resulting classifier on the vector *v_i_*. The final accuracy is computed as the average of the accuracies for all vectors. This technique fits our data set and its limited number of vectors. A more classical approach such as train-validation-test would have required us to use very small sets.

Importantly, we are also performing a second nested level of LOOCV to avoid any bias during the parametrization of the classifiers. This second level is described in section 3.3 and illustrated in *algorithm 1*.

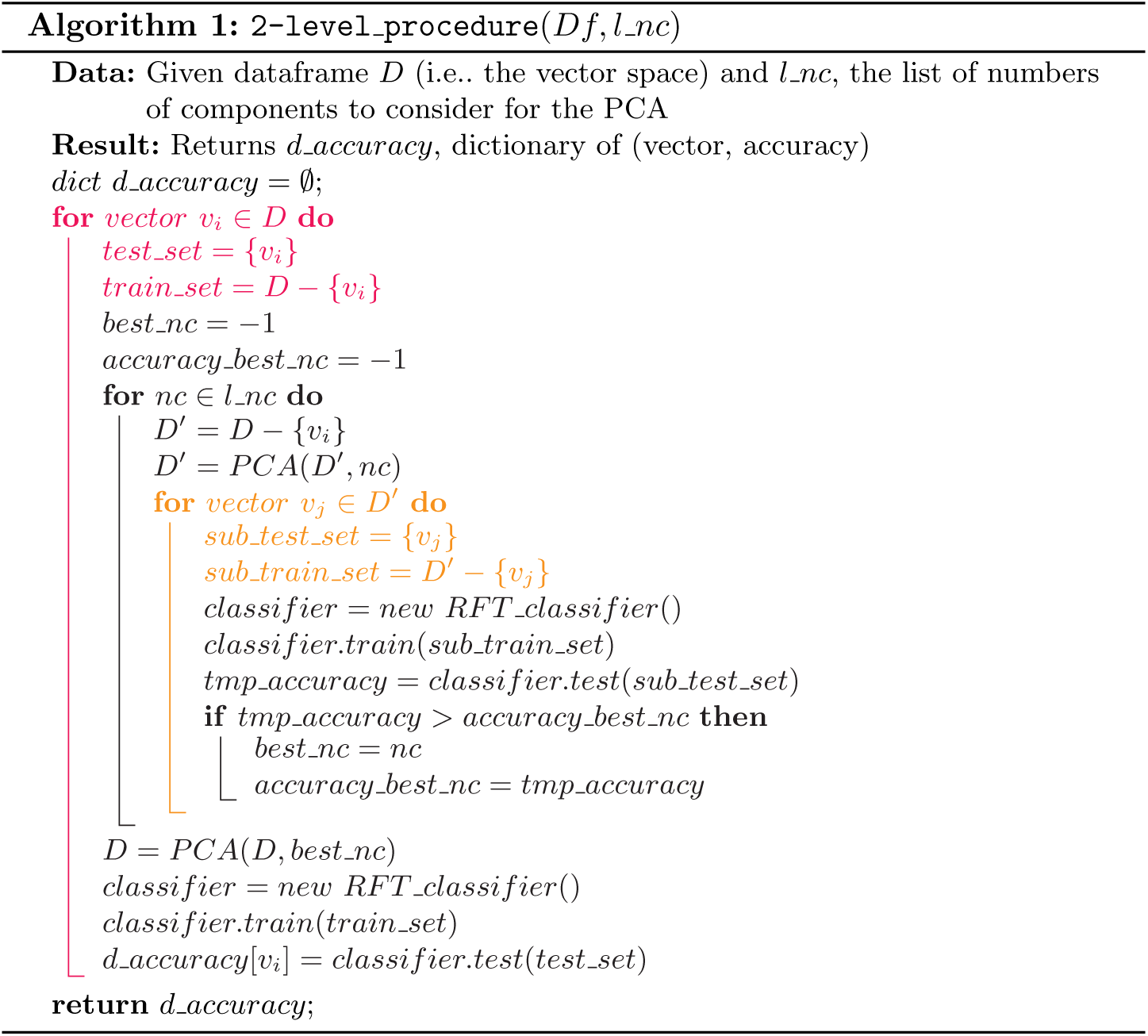

#### Principal Component Analysis (PCA)

Since the ratio *|vectors|/| features|* is poor in the dataset, it may hinder the accuracy of the classifiers. Principal component analysis (PCA) is a standard method to improve this ratio by reducing the number of dimensions. The PCA uses an orthogonal transformation to build a set of uncorrelated features (components) from the initial features with the objective of maximizing variance (i.e. minimizing the information lost by transforming).

The number of components to transform to is an important parameter that may influence the classifier accuracy. Performing preliminary tests to determine the best number would lead to a serious risk of overfitting. As a consequence we dynamically determined this number for each vector. The procedure is described in algorithm 1. From the first LOOCV, the set of vectors *V* has been split into a set of pairs of a training set *V′* = (*V* − {*v*_*i*_}) and a test set {*v*_*i*_}. For each pair, a second LOOCV is performed on *V′* leading to another set of pairs of a training set *V″* = (*V′* − {*v*_*j*_}) = (*V* − {*v*_*i*_, *v*_*j*_}) and a test set *{v*_*j}*_. Potential values for the number of components are tested and the one producing the best accuracy over *V′* is selected and used on *V* to predict the label of *v*_*i*_. As a consequence, the number of components to transform to is always selected independently from the test set.

Ideally, the set of potential values for the number of components would be 1, 2, …, |*V*|. However the computation time grows linearly with the number of values tested. As a consequence we decided to use a subset of 1, 2*, …, |V |* instead. Preliminary tests shows a light peak of performances at 8 10 components with a slight decrease before and after. As a consequence we tried all values from 1, 2*, …,* 20. We also added 0 (i.e. not performing a PCA).

#### Random Forest (RF) Classifier

We chose to use ensemble learning and more specifically Random Forest (RF) classifiers over other methods and classifiers because of some anticipated properties of the datasets. Indeed, the limitations of prediction tools are likely to generate noise which RF are relatively resilient to [21, p.596]. Moreover, the interactions we aimed at capturing were likely to be complex and the size of the training set to be limited. Since RF can capture complex interactions and are simple to train [21, p.587] compared to other classifiers [21, p.587] they appeared to be a fitting candidate.

Our implementation uses the python package Scikit-learn [22].

As the name suggests, Random Forest classifiers involve randomness. As a consequence we repeated the procedure and display distributions in order to counterbalance the variation of the predictions. Preliminary results show that the average accuracies of those distributions converge (10^*−*^4) within the first 500 runs. However we decided to double this value to add a comfortable security margin.

#### Dummy Classifier

A second classifier is trained in parallel to serve as a control. As the name “dummy” suggests, it is not an actual classifier but an heuristic randomly generating labels for the test set according to the probabilities distribution it extracted from the training set. As the dummy classifier is always trained and tested on the same sets as its RFT counterparts, it appears to be a suitable solution to produce a sound control while using LOOCV and using unbalanced labels. However, as all dummy classifiers produced extremely close performances, we decided to display only one of the dummy classifiers in each display instead of one per other classifier for the sake of clarity. Please note that the dummy classifier is unaffected by PCA as it does not consider the features.

#### Performance Metric for a Multi-label Dataset

The number of labels in our data sets prevents straightforward use of some classical displays such as ROC curves. A single prediction can indeed be, for instance and at the same time, both a false positive for a given label and a true negative for another. As a consequence we have *TPR* + *FPR* + *TNR* + *FNR* ≥ 1 ([True,False] [Positive,Negative] Rate) and plotting one ROC curve for each label offers little readability. As a consequence we instead chose to use displays based on accuracy (*Accuracy* = *P recision* = *|T rue P redictions|/|Predictions|*|).

### 3.4 Main Experiments

As described in section 3.2 and section 3.1, our work associates each ncRNA with three vectors of features ({Kmer,intaRNA,RNAup}) and a label. By doing so we produced three vector spaces which are suitable for machine learning. We consider that the ability of the classifiers to predict those labels reflects the said specificity. Therefore, the goal of the machine learning procedure described in section 3.3 is to assess the specificity of the sets of features regarding the labels.

Figure 2 displays the distribution of the accuracies of the classifiers for all three kinds of features with two or five labels. The combination of LOOCV (cf. 3.3) and the inherent randomness of RF classifiers (cf. 3.3) lead us to produce and display distributions of accuracies instead of a single value. Exact means (*µ*) and standard deviations (*σ*) values for all those distributions are displayed aside in Table 2. The best accuracies are obtained from the Kmer similarity scores with 86.3% of correct prediction with two labels and a standard variation of only 0.31%. Results obtained from scores predicted by RNAup are less accurate but are still very distinct from the control with no overlapping. However, results obtained from scores predicted by IntaRNA are significantly less accurate to the point that the distribution overlaps with the control. Results obtained with five labels display a similar hierarchy between Kmer, RNAup and IntaRNA with the addition of an expected global loss of accuracy. Indeed, the increased number of labels to predict makes the problem harder as shows the important drop of accuracy of the control. However, Kmer, RNAup and IntaRNA appear to all be more resilient than the control to this change.

**Fig. 1.**
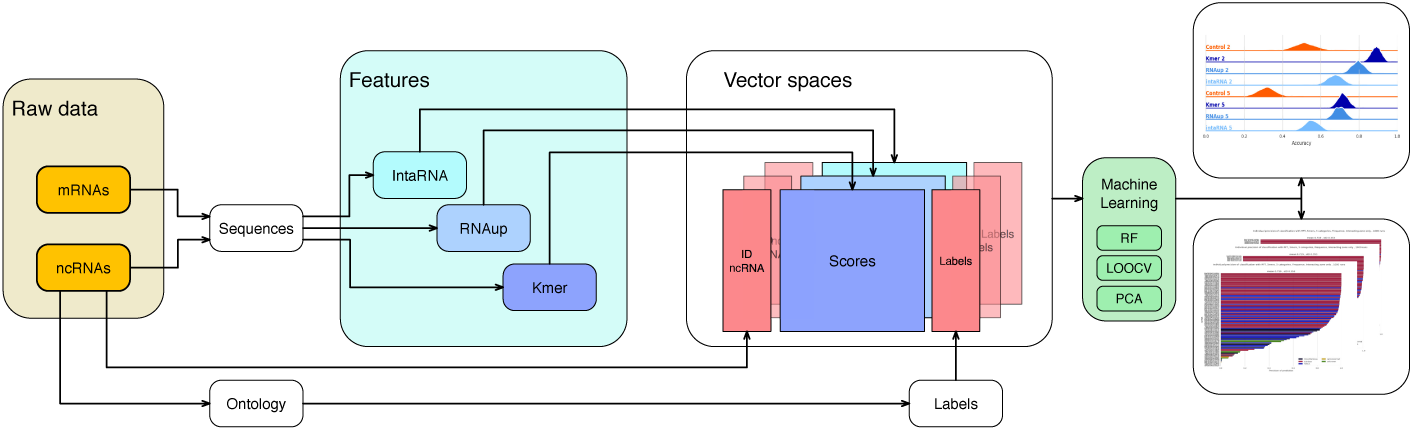
Illustration of our ensemble learning pipeline. The process starts from RNAs data in orange. Each ncRNA will be associated to a vector in the vector spaces and will be attributed a label according to its ontology. From either ncRNAs sequences (Kmer) or both ncRNA and mRNAs sequences (IntaRNA, RNAup), a set of scores will be computed and used as features. Machine learning is finally used to produce the results we are presenting from those vector spaces.

**Fig. 2.**
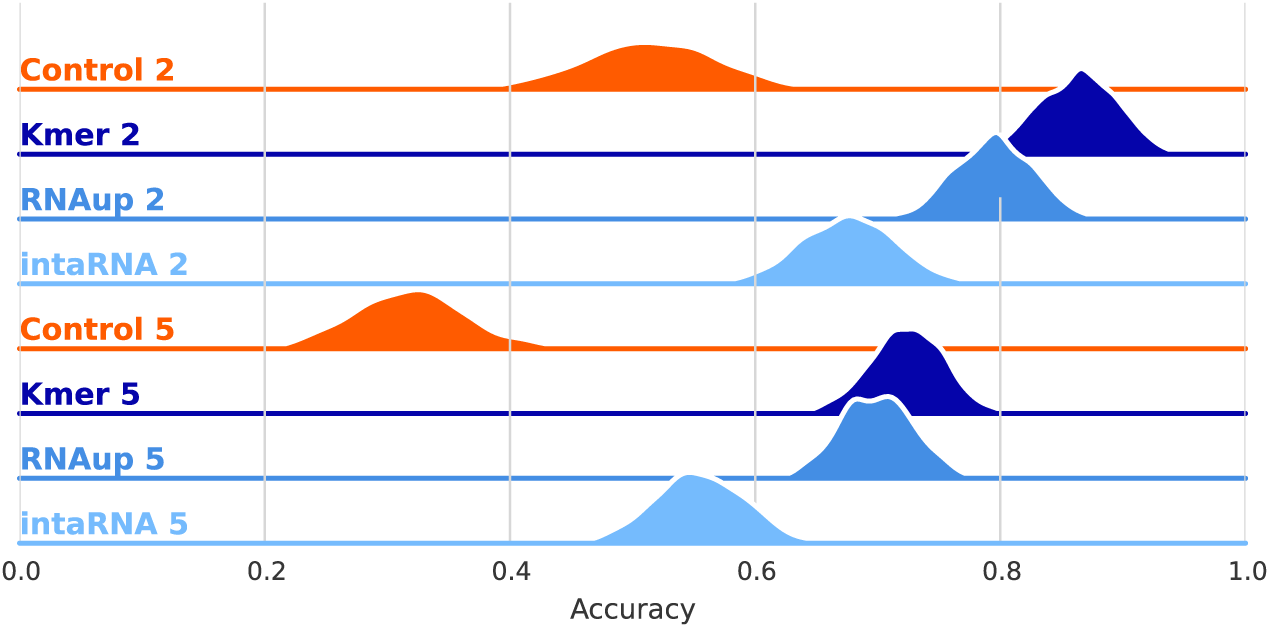
Distribution of accuracies of 1000 classifiers following specifications of section 3.3. Each row corresponds to either a set of features (*{*Kmer,RNAup,intaRNA*}*, cf. section 3.2) or the control (cf. section 3.3) associated with a number of labels (*{*2,5*}*, cf. section 3.1). Means and standard deviations for all distributions are displayed in Table 2.

**Table 2.** Means (*µ*) and standard deviations (*σ*) for all distributions displayed in Figure 2

|  | 2 labels |  | 5 labels |  |
| --- | --- | --- | --- | --- |
| | $\mu$ | $\sigma$ | $\mu$ | $\sigma$ |
| Control | 0.517 | 0.057 | 0.316 | 0.044 |
| Kmer | 0.863 | 0.031 | 0.724 | 0.03 |
| RNAup | 0.794 | 0.032 | 0.699 | 0.03 |
| intaRNA | 0.675 | 0.04 | 0.555 | 0.037 |

This first display suggests that the interaction profiles predicted by IntaRNA are significantly less specific than the ones predicted by RNAup. The interaction profiles predicted by RNAup also appear to be the closest to the ones produced from Kmer similarity scores and thus seem to give the most accurate account of the specificities of the sequences. This observation together with the difference between the two prediction tools described in section 3.2 suggest that probable interactions (i.e. the ones predicted by IntaRNA) are more inhibited than the potential optimal ones (i.e. the ones predicted by RNAup). This first observation is coherent with the influence of an evolutionary pressure as the inhibition of probable interactions would have a greater impact than the inhibition of potential optimal ones which are less likely to form.

Figure 3 is a different presentation of the results displayed in Figure 2. Raw results from the classifiers are unitary predictions (i.e. predictions of the label of one vector). We gathered those unitary predictions for each vector, thus producing an averaged accuracy for each of them. Figure 3 aims at highlithing variations inside the distribution displayed in Figure 2. Please note that each column corresponds now to a different set of features while the upper row displays the results with two labels and the lower row displays the results with five labels. Each line corresponds to a ncRNA, the length reflecting the accuracy of predictions made for this ncRNA label while the colour corresponds to its label. Please also note that lines are sorted by accuracies. As a consequence, the order varies in all of those six subgraphs.

**Fig. 3.**
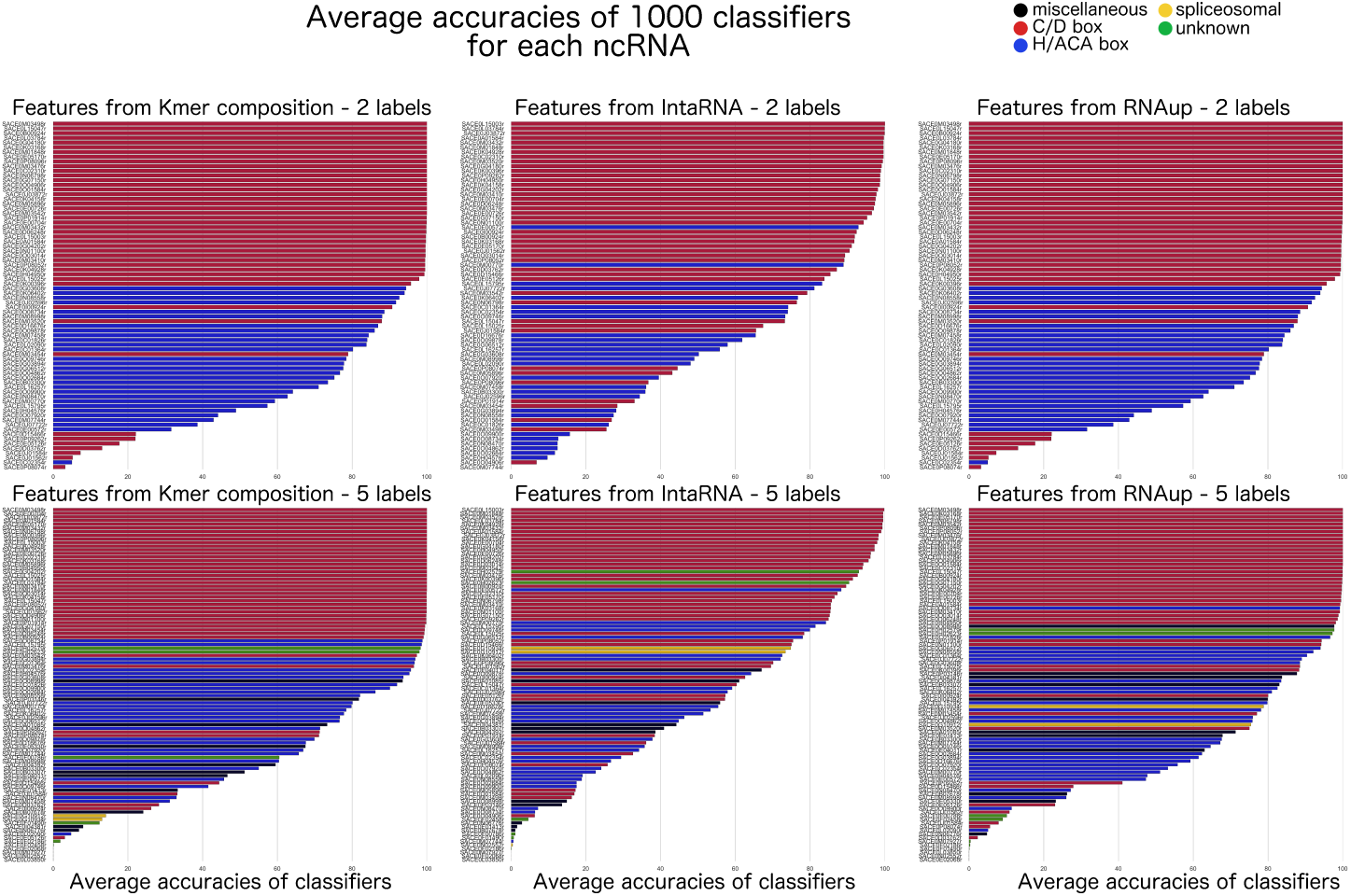
Average accuracies of classifiers for each vector (i.e. ncRNA) over 1000 tests. Each column corresponds to a set of features ({Kmer,RNAup,intaRNA}, cf. section 3.2). The first row displays the results with five labels and the second with two labels (cf. section 3.1). Each colored line in those 6 displays corresponds to a [vector/ncRNA]. The length of the line represents the averaged accuracy of the 1000 classifiers for the corresponding [vector/ncRNA]. The colour of the line corresponds to the label of the associated [vector/ncRNA]. Please note that [vectors/ncRNAs] are sorted according to the accuracy associated to them. As a consequence the order is different in all six graphs.

The drop of accuracy observed in Figure 2 between Kmer similarity scores, RNAup predicted scores and IntaRNA scores is also visible in Figure 3 as a more concave slope for better performing sets of features. However Figure 3 also displays variations of accuracies from one label to the other. C/D box RNAs (red) are the most noticeable group as those RNAs are, on average, extremely well-predicted with all features and either two and five labels. H/ACA box RNAs (blue), on the other hand, seem to be harder to predict from Kmer similarity scores or RNAup predicted scores than C/D box RNAs but show a dramatic drop of accuracy in predictions made from IntaRNA predicted scores. Predictions accuracies of the three remaining labels vary from a set of features to the other and even inside a label for a given set of features. We have been unable so far to determine if this was only due to a lesser number of vectors for those labels or to other parameters.

Results displayed in Figure 3 complement our previous observations as the Kmer,RNAup,IntaRNA hierarchy is still clearly observable. However, Figure 3 displays a phenomenon invisible in Figure 2: the variance in predictions accuracy between the label, especially regarding C/D box RNAs and H/ACA box RNAs. Indeed, predictions for C/D box RNAs (red) are always the most accurate while predictions for H/ACA box RNAs (blue) clearly fall behind. This variance goes from a limited difference (most predictions for H/ACA box RNAs are still above 80% accuracy in predictions from Kmer similarity scores with two labels, cf. top left graph) to a dramatic drop (predictions from IntaRNA predicted scores with two labels, cf. top right graph). Predictions accuracies of the three remaining labels vary from a set of features to the other and even inside a label for a given set of features. We have been unable so far to determine if this was only due to a lesser number of vectors for those labels or to other parameters. However, the predictions of the three remaining labels display accuracies similar to the ones of predictions for H/ACA box RNAs. Since the dataset contains more H/ACA box RNAs than the three other labels put together, this similarity stresses that H/ACA box RNAs are way harder to predict than C/D box RNAs. Further discussions of this difference of performances between labels require to first introduce Figure 4.

**Fig. 4.**
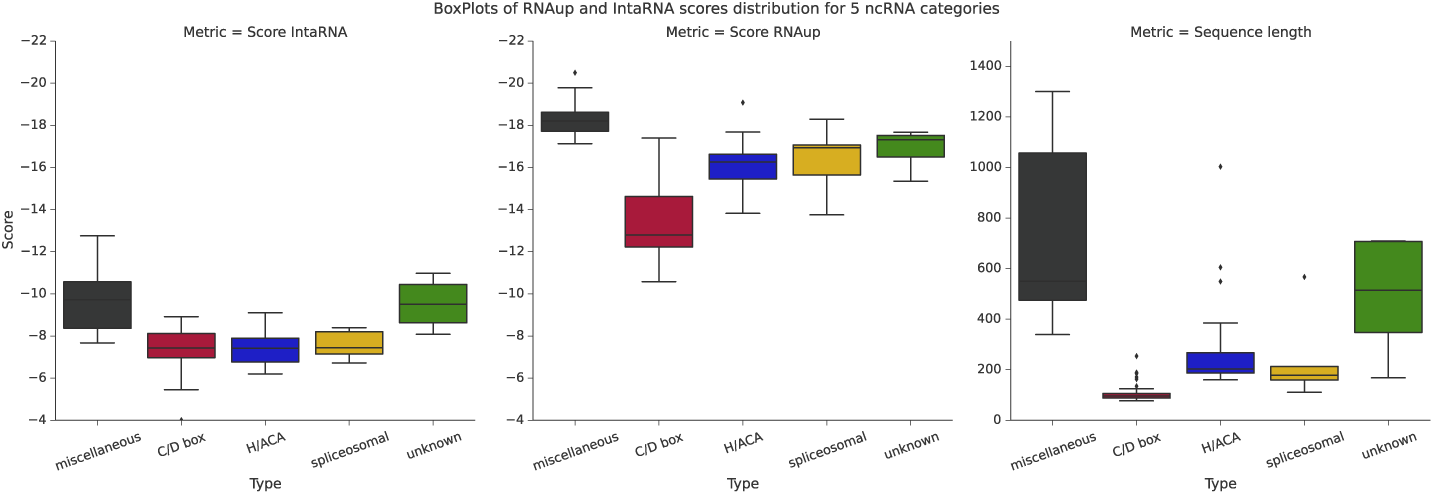
Normalized distributions of scores predicted by IntaRNA (left), scores predicted by RNAup (middle) and lengths of sequences (right) for each ncRNA label. Scores are in kcal/mol. Please note that scores from both IntaRNA and RNAup approximate a difference of entropy (*∆g*) and are therefore negative. A lower score thus suggests that the interaction is stronger.

In order to investigate the drop in accuracy between predictions made from scores predicted by RNAup and IntaRNA we plotted distributions of scores as box plots for each tool (left and middle) and for each ncRNA labels (colours). We also plotted the distributions of the lengths of ncRNAs sequences (right, please note that the influence of length results is discussed in section 3.5). The results are displayed in Figure 4. The colour code is the same as in Figure 2 and Figure 3. Figure 4 shows that RNAup is not only outputting stronger scores (entropy scores are negative cf. 3.1) but also preserves distinctions between the labels, especially between C/D box RNAs and H/ACA RNAs scores. This observation is coherent with the better performances of classifiers learning from the interactome predicted with RNAup. However the important drop in accuracies displayed in Figure 3 on scores predicted with IntaRNA with two labels shows that RF classifiers are able to capture variations (cf. Figure 2 and Figure 3) that the extremely similar distributions of those two labels in Figure 4 fail to display. This observation suggests that the global inhibition that is shown by the drop in the averages of both RNAup and IntaRNA scores is also a levelling phenomenon rather than a “linear” inhibition.

### 3.5 Additional Experiments

#### Impact of Boxes on Predictions

Among the five labels we are considering, two correspond to ncRNA classes defined by the presence of “boxes” in the sequence: C/D box snoRNAs and H/ACA box snoRNA. Those boxes are small and their consensus sequences are flexible (C: RUGAUGA, D: CUGA, H: ANANNA, ACA: ACA). Yet they might bias our results, especially those obtained from Kmer composition. To investigate this matter, we performed a brute force feature selection algorithm specific to random forest classifiers: the Boruta algorithm. This algorithm tests each feature, estimates its contribution to the classification and produces the list of features considered to be crucial for a given threshold of confidence (i.e. p-value, default = 0.05). Results show that only 50% or less of the critical features are compatible with a consensus sequence, even in the 2-label dataset (restrained to C/D box and H/ACA boxes snoRNAs only). This result is an upper bound since boxes are located while Kmer distributions ignore positions. As a consequence, the results displayed in Figure 2 and 3 cannot be produced only from Kmers capturing boxes.

#### Impact of Sequence Lengths on RNA-RNA Interaction Predictions

The third panel of Figure 4 displays the distributions of lengths of ncRNAs for all labels, each label being represented by a boxplot in its usual colour. Length distributions vary from a label to another with two visible groups of labels: *C/D box*, *H/ACA box* and *spliceosomal* labels (resp. red, blue and yellow) distributions are tightened around a relatively short length while *miscellaneous* and *unknown* labels (resp. black and green) present a wider distribution with overall longer sequences. The problem planted by lengths of mRNAs targets has been explored by *Umu et al.* [19, 20]. Their results show that the accuracy of prediction tools typically drops as the length of the target increases above 300nb. However, amongst the prediction tools tested, *IntaRNA* displays very little to no loss as the length of the target increases. On the contrary *RNAup* performances are significantly reduced. Cutting down the targets into subsequences of manageable length is not suited for this study as we need one score per {ncRNA,mRNA} pair. Moreover, we would like to propose to interpret this drop not only as a flaw of RNAup but as an illustration of the difference we described in 3.2. Yet the predictions scores for *miscellaneous* and *unknown* labels (resp. black and green) are to be treated with caution.

A second problem to consider is that the features we used are not independent of sequence lengths. Indeed, a longer sequence will contain more Kmers and Figure 4 suggests a partial correlation between scores and length. In order to investigate this issue we repeated the ensemble learning procedure with the length as the only feature. Results show that predictions using length are accurate (*µ*=0.856 and *σ*=0.012 with the 2-label dataset,*µ*=0.651 and *σ*=0.013 with the 5-label dataset) but are slightly outperformed by the ones trained on RNAup scores over the 5-label dataset and over both datasets by the ones trained on Kmer composition. Those results suggest that sequence lengths are specific to each labels but are not the only variation captured by the classifiers.

#### Overlapping of Predicted Interaction Zones with Observed Interaction Zones

We scanned the sequences of C/D box snoRNAs in the dataset looking for the consensus sequences. We excluded C/D box snoRNAs with ambiguous sites (i.e. more than one match with the consensus sequences of either box in the corresponding potential areas). We then looked for any intersection between the area interacting with rRNAs in observations (i.e. 3-rd to 11-th nucleotides upstream from D box) and the interaction zones predicted by RNAup. Amongst the interaction zones involving the 35 selected C/D box snoRNAs candidates, none overlapped with the observed interaction zones.

## 4 Conclusion

Our results enabled us to identify the signature of an evolutionary pressure against random interactions between ncRNAs and mRNAs in *Saccharomyces Cerevisae*. Presumably, as previously observed in prokaryotes and archaea, this phenomenon aims to increase the translation efficiency [10].

Although our data set includes various types of ncRNAs, the vast majority of them are snoRNAs. Our conclusions are therefore primarily applicable to snoRNAs, even if our data do not exclude that it could be generalized to other ncRNAs. Interestingly, the (old) age of the snoRNA family suggests that it coud be the trace of a fundamental biological process used by primitive microorganisms. The absence (to our knowledge) of experimental evidences of snoRNA-mRNA interactions in unicellular eukaryotes tends to support our conclusions. By contrast, the existence of known interactions between orphan snoRNAs and mRNAs in human or mice [4, 5] opens a legitimate debate about the necessity and specificity of such mechanisms in animals.

